# The unequal functional redundancy of Arabidopsis *INCURVATA11* and *CUPULIFORMIS2* is not dependent on genetic background

**DOI:** 10.1101/2023.04.20.537354

**Authors:** Riad Nadi, Lucía Juan-Vicente, Eduardo Mateo-Bonmatí, José Luis Micol

**Affiliations:** Instituto de Bioingeniería, Universidad Miguel Hernández, Campus de Elche, 03202 Elche, Spain; Current address: Centro de Biotecnología y Genómica de Plantas (CBGP), Universidad Politécnica de Madrid (UPM) – Instituto Nacional de Investigación y Tecnología Agraria y Alimentaria (INIA)/CSIC, 28223 Pozuelo de Alarcón, Madrid, Spain

**Keywords:** epigenetic machinery, *Arabidopsis thaliana*, *ICU11*, 2OGD, *CUPULIFORMIS*, CRISPR/Cas9.

## Abstract

The paralogous genes *INCURVATA11* (*ICU11*) and *CUPULIFORMIS2* (*CP2*) encode components of the epigenetic machinery in Arabidopsis and belong to the 2-oxoglutarate and Fe (II)-dependent dioxygenase superfamily. We previously inferred unequal functional redundancy between *ICU11* and *CP2* from a study of the synergistic phenotypes of the double mutant and sesquimutant combinations of *icu11* and *cp2* mutations, although they represented mixed genetic backgrounds. To avoid potential confounding effects arising from different genetic backgrounds, we generated the *icu11-5* and *icu11-6* mutants via CRISPR/Cas genome editing in the Col-0 background and crossed them to *cp2* mutants in Col-0. The resulting mutants exhibited a postembryonic-lethal phenotype reminiscent of strong *embryonic flower* (*emf*) mutants. Double mutants involving *icu11-5* and mutations affecting epigenetic machinery components displayed synergistic phenotypes, whereas *cp2-3* did not besides *icu11-5*. Our results confirmed the unequal functional redundancy between *ICU11* and *CP2* and demonstrated that it is not allele or genetic background specific. An increase in sucrose content in the culture medium partially rescued the post-germinative lethality of *icu11 cp2* double mutants and sesquimutants, facilitating the study of their morphological phenotypes throughout their life cycle, which include floral organ homeotic transformations. We thus established that the *ICU11-CP2* module is required for proper flower organ identity.

## INTRODUCTION

In *Arabidopsis thaliana* (hereafter, Arabidopsis), as in many other model species, the genetic dissection of biological phenomena typically involves the isolation and genetic analysis of mutants (Nüsslein-Volhard and Wieschaus, 1980; Jürgens et al., 1991; Koornneef et al., 1991; Wilkins, 1992; Haffter et al., 1996; Berná et al., 1999). The choice of wild-type strain to be mutagenized is a key step in this endeavor, as mutant phenotypes clearly distinguishable from the wild type will not be produced for some genes in some genetic backgrounds (Lee et al., 1994; Koornneef et al., 2004; Chandler et al., 2013; Leng et al., 2022).

In addition, comparative analysis of the morphological, physiological, and molecular phenotypes of double or higher-order mutant combinations obtained by crossing single mutants is not always straightforward, given that different genetic backgrounds sometimes need to be mixed because of mutant availability in distinct strains (Huq and Quail, 2002; Scortecci et al., 2003; Clerkx et al., 2004). Indeed, phenotypes may be strongly influenced by modifiers present in the genomes of wild-type strains subjected to mutagenesis (Fernando et al., 2018). One strategy to partially overcome this problem is to first introgress each mutation of interest into an adequate genetic background, but this approach is time consuming and often leaves traces of the donor background (Rustérucci et al., 2001; Mouchel et al., 2004; Yoo et al., 2007; Zikherman et al., 2009; Kradolfer et al., 2013). An alternative approach is now accessible via clustered regularly interspaced short palindromic repeats (CRISPR)/CRISPR-associated nuclease (Cas)-mediated genome editing, which allows the relatively rapid isolation of single or multiple mutants in the same genetic background. CRISPR/Cas9 is now the preferred choice for directed mutagenesis due to its high specificity, efficiency, and simplicity (Jinek et al., 2012; Cong et al., 2013; Gaj et al., 2013; Jia et al., 2016; Zhao et al., 2016; Wang et al., 2022).

The Arabidopsis paralogous epigenetic factors INCURVATA11 (ICU11) and CUPULIFORMIS2 (CP2; Mateo-Bonmatí et al., 2018) belong to one of the largest known protein superfamilies, the 2-oxoglutarate and Fe (II)-dependent dioxygenases (2OGDs), which is represented by about 150 members in plants (Kawai et al., 2014; Martinez and Hausinger, 2015; Nadi et al., 2018). These proteins catalyze oxidation reactions using 2-oxoglutarate (also called α-ketoglutarate) and molecular oxygen as cosubstrates, and ferrous iron (Fe^2+^) as a cofactor (Islam et al., 2018). ICU11 is a POLYCOMB REPRESSIVE COMPLEX 2 (PRC2) accessory protein likely involved in removing the active histone mark H3K36me3 (trimethylation of lysine 36 of histone H3; Bloomer et al., 2020).

We previously described unequal functional redundancy between *ICU11* and *CP2* (Mateo-Bonmatí et al., 2018). The *icu11-1* and *icu11-2* mutant alleles in the S96 and Wassilewskija-2 (Ws-2) genetic backgrounds, respectively, showed mild but pleiotropic phenotypic defects, such as early flowering and curled (hyponastic) rosette leaves, while the *cp2-1*, *cp2-2*, and *cp2-3* alleles in Columbia-0 (Col-0) were indistinguishable from their wild type. Notably, double mutant combinations between the *icu11* null alleles and the hypomorphic *cp2* alleles *cp2-1* and *cp2-2* skipped the vegetative phase and flowered immediately after germination, producing aberrant and sterile embryonic flowers. Double mutants with the null *cp2-3* allele were not obtained. The reciprocal sesquimutants *ICU11/icu11-1*;*cp2-3*/*cp2-3* and *icu11-1*/*icu11-1*;*CP2*/*cp2-3*, each only harboring one functional gene copy out of four, were not equivalent: while one copy of *ICU11* was sufficient to obtain plants that were phenotypically wild type, a single copy of *CP2* was not, with this sesquimutant producing lethal embryonic flowers (Mateo-Bonmatí et al., 2018).

Here, we obtained by CRISPR/Cas9-mediated gene editing alleles of *ICU11* in the Col-0 and S96 backgrounds. They had differing phenotypes as single mutants but apparently identical genetic interactions with *cp2* alleles in double mutant combinations. We therefore provide evidence that the lethal postembryonic phenotype of the *icu11 cp2* double mutants and sesquimutants is not specific to the allele or the genetic background. We also discovered that this seedling lethality can be circumvented by increasing the sucrose content of the growth medium, which in turn allowed us to obtain evidence of the requirement of the *ICU11-CP2* module for proper flower organ identity.

## METHODS

### Plant material, culture conditions, and crosses

Unless otherwise stated, all *Arabidopsis thaliana* (L.) Heynh. plants studied in this work were homozygous for the mutations indicated. The Nottingham Arabidopsis Stock Centre (NASC) provided seeds for the wild-type accessions Columbia-0 (Col-0; N1092), S96 (N914), and Wassilewskija-2 (Ws-2; N1601), as well as the following mutants: *icu11-1* (N242) in the S96 background; *curly leaf-2* (*clf-2*; N8853) in the Landsberg *erecta* (L*er*) background; *arabidopsis trithorax1-2* (*atx1-2*; N649002), *arabidopsis trithorax-related protein 5* (*atxr5*; N630607), *atxr6* (N866134), *atxr7-1* (N667600), *cp2-1* (N861581), *cp2-2* (N828642), *cp2-3* (N826626), *demeter-like 2-3* (*dml2-3*; N631712), *dml3-1* (N556440), *dna methyltransferase-2-2* (*dnmt2-2*; N836854), *domains rearranged methylase 1-2* (*drm1-2*; N521316), *drm2-2* (N650863), *histone acetyltransferase of the cbp family 1-3* (*hac1-3*; N580380), *histone acetyltransferase of the myst family 1-1* (*ham1-1*; N655396), *methyltransferase 1-4* (*met1-4*; N836155), *repressor of silencing 1-4* (*ros1-4*; N682295), *ros3-2* (N522363), *terminal flower2-2* (*tfl2-2*; N3797), and *embryonic flower2-3* (*emf2-3*; N16240) in the Col-0 background; *icu2-1* (N329) and *fasciata1-1* (*fas1-1*; N265) in the Enkheim2 (En-2) background; *methyl-cpg-binding domain10-1* (*mbd10-1*; N872244) and *variant in methylation 3-2* (*vim3-2*; N804664) in the Col-3 background; and *histone deacetylase 6-6* (*hda6-6*; N66153) and *hda6-7* (N66154) in the Col background. Seeds for *early bolting in short days-1* (*ebs-1*, in the L*er* background; Piñeiro et al., 2003) were provided by Manuel Piñeiro (CBGP, UPM-INIA-CSIC, Madrid, Spain), those of *gigantea supressor5* (*gis5*, in the Col-0 background; Iglesias et al., 2015) by Pablo D. Cerdán (Fundación Instituto Leloir, IIBBA-CONICET, Buenos Aires, Argentina), and those of *icu11-2* (in the Ws-2 background) by the Versailles Arabidopsis Stock Center (Brunaud et al., 2002). The presence and positions of all T-DNA insertions were confirmed by PCR amplification using gene-specific primers, with the LbB1.3 and LB1 primers used for the SALK and SAIL T-DNA insertions, respectively (Supplementary Table S1).

Unless otherwise stated, all seeds were surface sterilized, plated onto 140-mm (diameter) Petri dishes containing 100 ml half-strength Murashige and Skoog (MS) plant agar medium with 1% (w/v) sucrose at 20 ± 1°C, 60-70% relative humidity, and continuous illumination at ∼75 µmol m^−2^ s^−1^, as previously described (Ponce et al., 1998). Crosses were performed as previously described (Quesada et al., 2000).

### Plant morphology and pollen staining

Photographs showing morphology were taken with a Nikon SMZ1500 stereomicroscope equipped with a Nikon DXM1200F digital camera. Pollen grains were stained with Alexander red solution for 5–10 min before observation and photographed using a Leica DMRB microscope equipped with a Nikon DXM1200 digital camera.

### Gene constructs and plant transformation

The pKI1.1R-ICU11_sgRNA1 plasmid was constructed as described by Tsutsui and Higashiyama (2017). In brief, the pKI1.1R plasmid (Addgene) was linearized by restriction digest with *Aar*I (Thermo Fisher Scientific), and treated with FastAP alkaline phosphatase (Thermo Fisher Scientific). The ICU11_sgRNA1_F/R oligonucleotides were phosphorylated using T4 polynucleotide kinase (New England Biolabs) and hybridized in a thermal cycler (Bio-Rad Laboratories T100). The ligation reaction was performed with T4 DNA ligase (Thermo Fisher Scientific), and the ligation product was transformed into chemically competent *Escherichia coli* DH5α cells using the heat-shock method. Plasmid and insert integrity were verified by Sanger sequencing using an Applied Biosystems 3500 Genetic Analyzer (Thermo Fisher Scientific). Putative off-targets were identified using the default parameters of the ChopChop tool (Labun et al., 2019). The pKI1.1R-ICU11_sgRNA1 plasmid was mobilized into *Agrobacterium tumefaciens* GV3101 (C58C1 Rif^R^) cells, which were used to transform Arabidopsis S96 and Col-0 plants via the floral dip method (Clough and Bent, 1998). T_1_ Arabidopsis transgenic plants were selected on plates with MS medium containing 15 mg l^−1^ hygromycin B (Thermo Fisher Scientific).

### Flowering time analysis

Flowering time was determined based on the total number of rosette leaves (counted when internode elongation was visible) and the number of days to bolting (Bouveret et al., 2006). To determine flowering time, all plants were grown on MS medium for five days and transferred to soil (a 2:2:1 mixture of perlite, vermiculite and sphagnum moss) in individual pots in a TC30 growth chamber (Conviron).

### Accession numbers

Sequence data from this article can be found at The Arabidopsis Information Resource (http://www.arabidopsis.org) under the following accession numbers: *ICU11* (At1g22950), *CP2* (At3g18210), *EBS* (At4g22140), *FAS1* (At1g65470), *GIS5* (At5g63960), *ICU2* (At5g67100), *CLF* (At2g23380), *TFL2* (At5g17690), *EMF2* (At5g51230), *DML2* (At3g10010), *DML3* (At4g34060), *DNMT2* (At5g25480), *DRM1* (At5g15380), *DRM2* (At5g14620), *MBD10* (At1g15340), *MET1* (At5g49160), *ROS1* (At2g36490), *ROS3* (At5g58130), *VIM3* (At5g39550), *ATX1* (At2g31650), *ATXR5* (At5g09790), *ATXR6* (At5g24330), *ATXR7* (At5g42400), *HAC1* (At1g79000), *HDA6* (At5g63110), and *HAM1* (At5g64610).

## RESULTS

### Isolation of novel *icu11* mutants in a Col-0 background after CRISPR/Cas9 mutagenesis

We previously characterized two loss-of-function *icu11* alleles: *icu11-1* and *icu11-2* (Mateo-Bonmatí et al., 2018). A third allele, *icu11-3*, was identified in an *Ac/Ds* transposon-tagging mutagenesis screen (Bancroft et al., 1993; Bloomer et al., 2020). Although these three mutants appear to be null alleles (Figure 1A), the hyponasty of *icu11-1* leaves is stronger than that of the *icu11-2* and *icu11-3* alleles, which is likely due to their different genetic backgrounds (S96, Ws-2, and L*er*, respectively). Different or hybrid genetic backgrounds may not facilitate proper comparisons of the morphological and molecular phenotypes of single and multiple mutants (Page and Grossniklaus, 2002; Chandler et al., 2013; Taylor and Ehrenreich, 2015); hence, as all *cp2* alleles were in the Col-0 background but there were no available *icu11* alleles in this background, we subjected Col-0 plants to CRISPR/Cas9 mutagenesis targeting *ICU11*. We also mutagenized S96 plants as a control. Accordingly, we designed a 20-nt single guide RNA (sgRNA) that targets the first exon of the *ICU11* gene, with a predicted targeting efficiency of 49.7% (Fig. 1A, B; Supplementary Fig. S1A). We obtained seven chimeric T_1_ plants, from which we isolated four independent homozygous lines: *icu11-4* and *icu11-7* mutants in the S96 background, carrying deletions of 4 bp and 1 bp, respectively, and *icu11-5* and *icu11-6* in the Col-0 background, carrying a 1-bp insertion and a 1-bp deletion, respectively (Fig. 1A; Supplementary Fig. S1B). All these mutations are predicted to cause frameshifts that introduce premature stop codons, producing truncated proteins of only 86 (*icu11-4*), 44 (*icu11-5*), or 87 (*icu11-6* and *icu11-7*) amino acids, instead of the 397 amino acids of wild-type ICU11 (Supplementary Fig. S1C). We sequenced *Cas9*-free lines by Sanger sequencing to examine the two most likely off-targets, both of which presented four mismatches with at least one mismatch located in the 5 bp adjacent to the protospacer adjacent motif (PAM) of our sgRNA (Supplementary Fig. S2A); neither off-target was mutated in subsequent generations of our mutant lines (Supplementary Fig. S2B).

**Figure 1.**
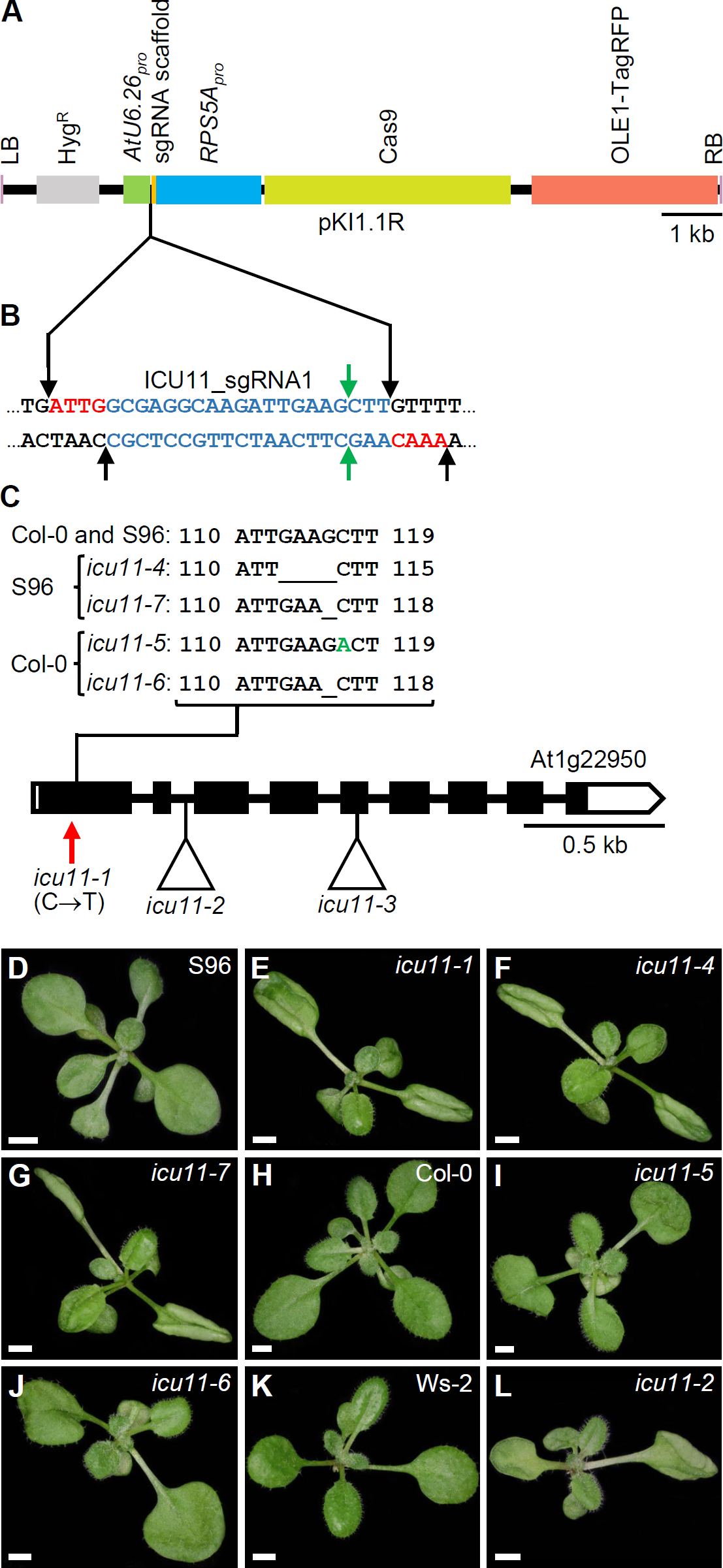
Molecular nature and morphological phenotypes of the *icu11* mutations obtained in this work. (A) Diagram of the T-DNA fragment of the pKIR1.1R vector that is integrated into the plant genome. Structural features are represented as boxes: LB and RB, left and right T-DNA borders (pink); *Hyg^R^*, hygromycin resistance gene (gray); *AtU6.26pro*, promoter of *U6 SMALL NUCLEOLAR RNA26* (green); sgRNA scaffold, the sequence that serves as a binding site for *Streptococcus pyogenes* Cas9 protein (orange); *RPS5Apro*, promoter of *RIBOSOMAL PROTEIN 5A* (blue); Cas9, CRISPR-associated protein 9 (pale green); *OLE1-TagRFP*, a translational fusion of OLEOSIN 1, the most abundant oleosin in Arabidopsis seeds, and the red fluorescent protein (RFP; red). Between the *AtU6.26* promoter and the sgRNA scaffold, there are two restriction sites for the type IIS *Aar*I restriction enzyme. (B) Nucleotide sequence of the sgRNA scaffold (black) and the *ICU11* sgRNA1 (blue) with four-nucleotide overhangs used for cloning (red). Black and green arrows indicate the *Aar*I restriction sites and predicted Cas9 breakpoints, respectively. (C) Schematic representation of the structure of the *ICU11* gene with indication of the nature and positions of *icu11* mutations. White and black boxes represent untranslated and coding regions of exons, respectively; lines represent introns. A red vertical arrow indicates the *icu11-1* point mutation, and triangles indicate the *icu11-2* T-DNA and *icu11-3 Ds* insertions (not studied in this work). The sequences of the *icu11-4*, *icu11-6*, and *icu11-7* deletions and the *icu11-5* insertion (+1 bp, in green) are also shown. (D–L) Rosettes of the S96 (D), Col-0 (H), and Ws-2 (K) wild-type accessions, and the *icu11-1* (E), *icu11-4* (F), *icu11-7* (G), *icu11-5* (I), *icu11-6* (J), and *icu11-2* (L) single mutants. Photographs were taken 15 days after stratification (das). Scale bars, 2 mm.

### The seedling-lethal phenotype of the *icu11 cp2* double mutants is independent of genetic background

As expected from their S96 background, the *icu11-4* and *icu11-7* mutants exhibited a morphological phenotype indistinguishable from that of *icu11-1* (Fig. 1C–F). The *icu11-5* and *icu11-6* mutants showed leaf hyponasty to a lesser extent (Fig. 1G–I), similar to that of *icu11-2* (Fig. 1J, K) and *icu11-3* (Bloomer et al., 2020). In addition, *icu11-5* and *icu11-6* shared other phenotypic traits with the other *icu11* mutants, such as cotyledon epinasty (Fig. 1C–K) and early flowering (Fig. 2A, B). We observed significantly more unfertilized ovules per half silique in the *icu11-4*, *icu11-5*, *icu11-6* and *icu11-7* mutants compared to their respective wild types (Fig. 2C–L). Taken together, these observations indicate that our new *icu11* mutants are phenotypically similar to previously reported mutants, with only minor differences caused by their genetic background, which are particularly visible in their rosette leaf morphology.

**Figure 2.**
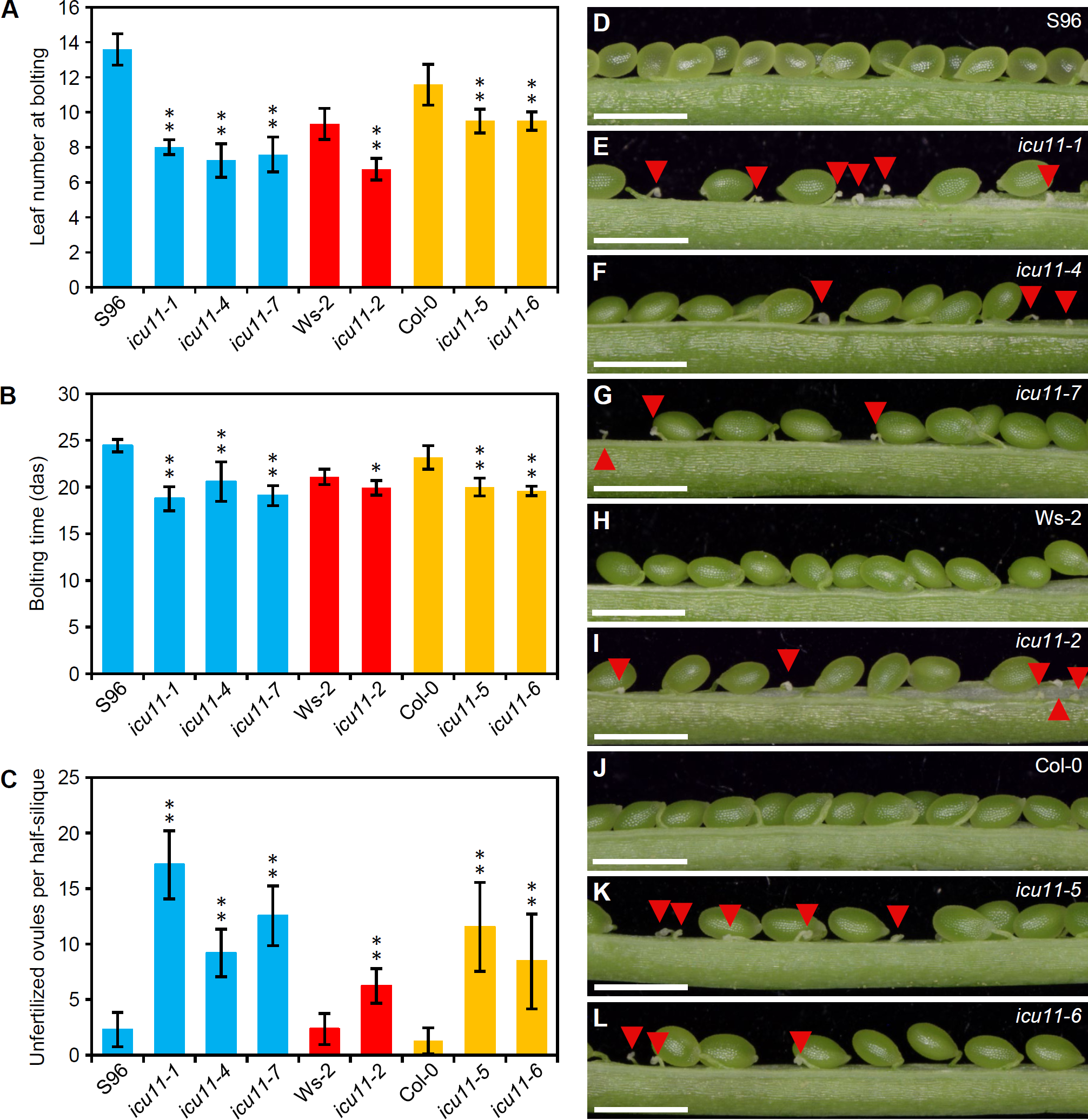
Flowering and reproductive phenotypes of the *icu11-5* and *icu11-6* mutants. (A–B) Flowering time of S96, *icu11-1*, *icu11-4*, *icu11-7*, Ws-2, *icu11-2*, Col-0, *icu11-5*, and *icu11-6* plants, expressed as leaf number at bolting (A) and number of days to bolting (B). (C) Number of unfertilized ovules per half silique for the indicated genotypes. Data are means ± standard deviation. Asterisks indicate values significantly different from the corresponding wild type in a Mann-Whitney *U* test (\**P* < 0.01 and \*\**P* < 0.001). Blue, red, and yellow bars indicate that the genotypes are in the S96, Ws-2, and Col-0 backgrounds, respectively. (D–L) Dissected fully elongated siliques for the indicated genotypes. Red arrowheads indicate unfertilized ovules. Photographs were taken 45 das. Scale bars, 1 mm. All siliques in C–L were collected from plants grown simultaneously within the same growth chamber. Pictures in D, E, H, I and J are similar to those that we published in the Supplemental Figure 2 (A, C, B, D and G) of Mateo-Bonmatí et al. (2018), respectively. Siliques of S96, *icu11-1*, Ws-2, *icu11-2* and Col-0 are included here to allow comparison with *icu11-4*, *icu11-7*, *icu11-5* and *icu11-6*. The numbers shown in C for S96, *icu11-1*, Ws-2, *icu11-2* and Col-0 have been obtained independently of those of Supplemental Figure 2L of Mateo-Bonmatí et al. (2018).

Although the morphological phenotypes caused by the *icu11* alleles are relatively mild, and the *cp2* null mutants are indistinguishable from wild type, the phenotypes of *icu11 cp2* double mutants are synergistic: they are seedling lethal, as might be expected for the genetic combination of mutations in two close paralogs with a high degree of functional redundancy (Ohno, 1970; Nowak et al., 1997; Cusack et al., 2021). Our previously obtained *icu11 cp2* double mutants had hybrid genetic backgrounds, which prevented a clear conclusion as to the embryonic lethality presented by these double mutants. Here, we thus crossed *icu11-5* and *icu11-6* with the *cp2-1* and *cp2-2* hypomorphic alleles of *CP2*, and with the *cp2-3* null allele, all of which are in the Col-0 genetic background. The double homozygous mutant combinations between *icu11-5* or *icu11-6* and *cp2-1* or *cp2-2* exhibited an embryonic-flowering seedling-lethal phenotype, as did the *icu11/icu11;CP2/cp2-3* sesquimutant (Fig. 3). We obtained no *icu11-5 cp2-3* or *icu11-6 cp2-3* double mutants, as was previously published for *icu11-1 cp2-3* (Mateo-Bonmatí et al., 2018). The *ICU11/icu11;cp2-3/cp2-3* sesquimutants were indistinguishable from the wild type, as were those we previously published in hybrid genetic backgrounds (Fig. 3I, N).

**Figure 3.**
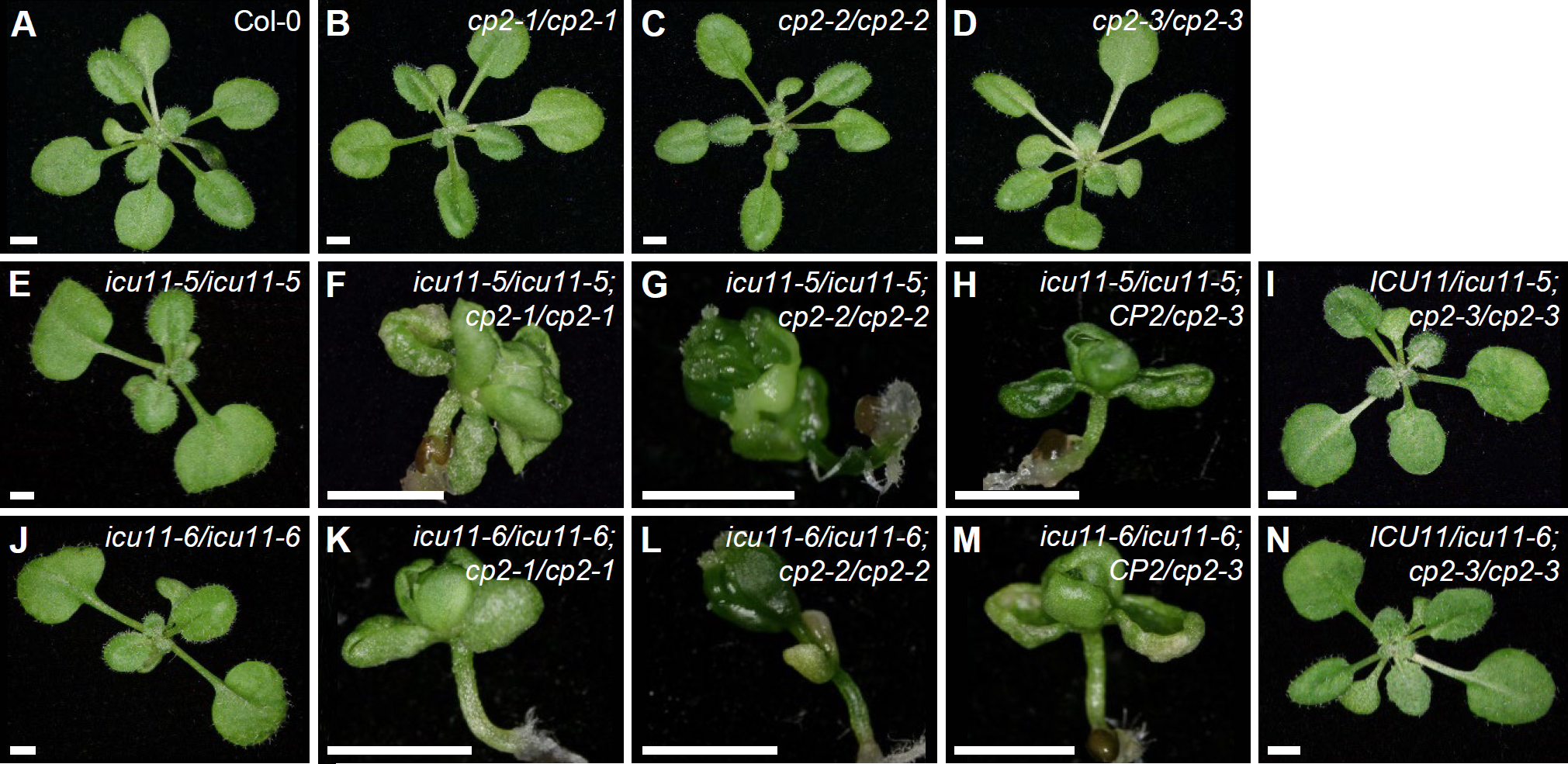
Genetic interactions between the loss-of-function *icu11* and *cp2* alleles in the Col-0 genetic background. *icu11-5*, *icu11-6*, and *cp2-3* are null alleles, while *cp2-1* and *cp2-2* are hypomorphic. Rosettes of the wild-type Col-0 (A); the homozygous single mutants *cp2-1* (B), *cp2-2* (C), *cp2-3* (D), *icu11-5* (E), and *icu11-6* (J); the double mutants *icu11-5 cp2-1* (F), *icu11-5 cp2-2* (G), *icu11-6 cp2-1* (K), and *icu11-6 cp2-2* (L); and the sesquimutants *icu11-5/icu11-5*;*CP2/cp2-3* (H), *ICU11/icu11-5*;*cp2-3/cp2-3* (I), *icu11-6/icu11-6*;*CP2/cp2-3* (M) and *ICU11/icu11-6*;*cp2-3/cp2-3* (N). Photographs were taken 16 das. Scale bars, 2 mm.

### *icu11-5*, but not *cp2-3*, genetically interacts with loss-of-function alleles of genes encoding PRC2 core components and accessory proteins

Synergistic phenotypes visualized in double mutants shed light on functional relationships between genes (Pérez-Pérez et al., 2009), including those encoding components of the epigenetic machinery of Arabidopsis. For example, PWWP-DOMAIN INTERACTOR OF POLYCOMB1 (PWO1) is a histone reader that recruits PcG proteins (Hohenstatt et al., 2018), and BLISTER (BLI) is a PRC2 interactor and a regulator of stress-responsive genes (Kleinmanns et al., 2017). Both *pwo1* and *bli* loss-of-function mutations display synergistic phenotypes when combined with strong mutant alleles of *CURLY LEAF* (*CLF*), which encodes a PRC2 core component responsible for the deposition of H3K27me3 repressive marks (Goodrich et al., 1997; Schatlowski et al., 2010).

With the aim to expand the spectrum of genes demonstrated to genetically interact with *ICU11*, we crossed *icu11-1* to loss-of-function mutants of 17 genes encoding components of the epigenetic machinery that include proteins involved in DNA or histone methylation, acetylation, or deacetylation (Pikaard and Mittelsten Scheid, 2014). We identified 18 double mutants with additive phenotypes (Supplementary Table S2). Also we obtained double mutant combinations of *icu11-5* with loss-of-function alleles of *CLF*, *LIKE HETEROCHROMATIN PROTEIN 1* (*LHP1*; also named *TFL2*), *FAS1*, *EBS*, *GIS5*, and *ICU2,* previously found to genetically interact with *icu11-1* (Mateo-Bonmatí et al., 2018). FAS1 is a component of the Chromatin Assembly Factor 1 (CAF-1) complex that promotes the deposition of histone H3 and H4 at newly synthesized DNA during replication (Schönrock et al., 2006). EBS is an H3K27me3 and H3K4me3 reader that regulates the floral phase transition (Piñeiro et al., 2003; López-González et al., 2014; Yang et al., 2018). GIS5 and ICU2 are the catalytic subunits of DNA polymerase δ and α, respectively (Barrero et al., 2007; Iglesias et al., 2015). The double mutant combinations of *icu11-5* with *clf-2*, *ebs-1*, *gis5*, *icu2-1*, *tfl2-2*, or *fas1-1* exhibited strong synergistic phenotypes consisting of dwarf rosettes; extreme leaf hyponasty in the cases of *gis5*, *icu2-1*, and *fas1-1*; and some degree of anthocyanin accumulation in the *icu11-5 tfl2-2* rosette center (Fig. 4). The phenotypes of the double mutant combinations of *icu11-5* with *clf-2*, *gis5,* and *icu2-1* were similar to those previously reported using *icu11-1*; however, those involving *tfl2-2*, *ebs-1*, and *fas1-1* were milder in combination with *icu11-5* (Fig. 4L–N) than with *icu11-1* (Mateo-Bonmatí et al., 2018).

**Figure 4.**
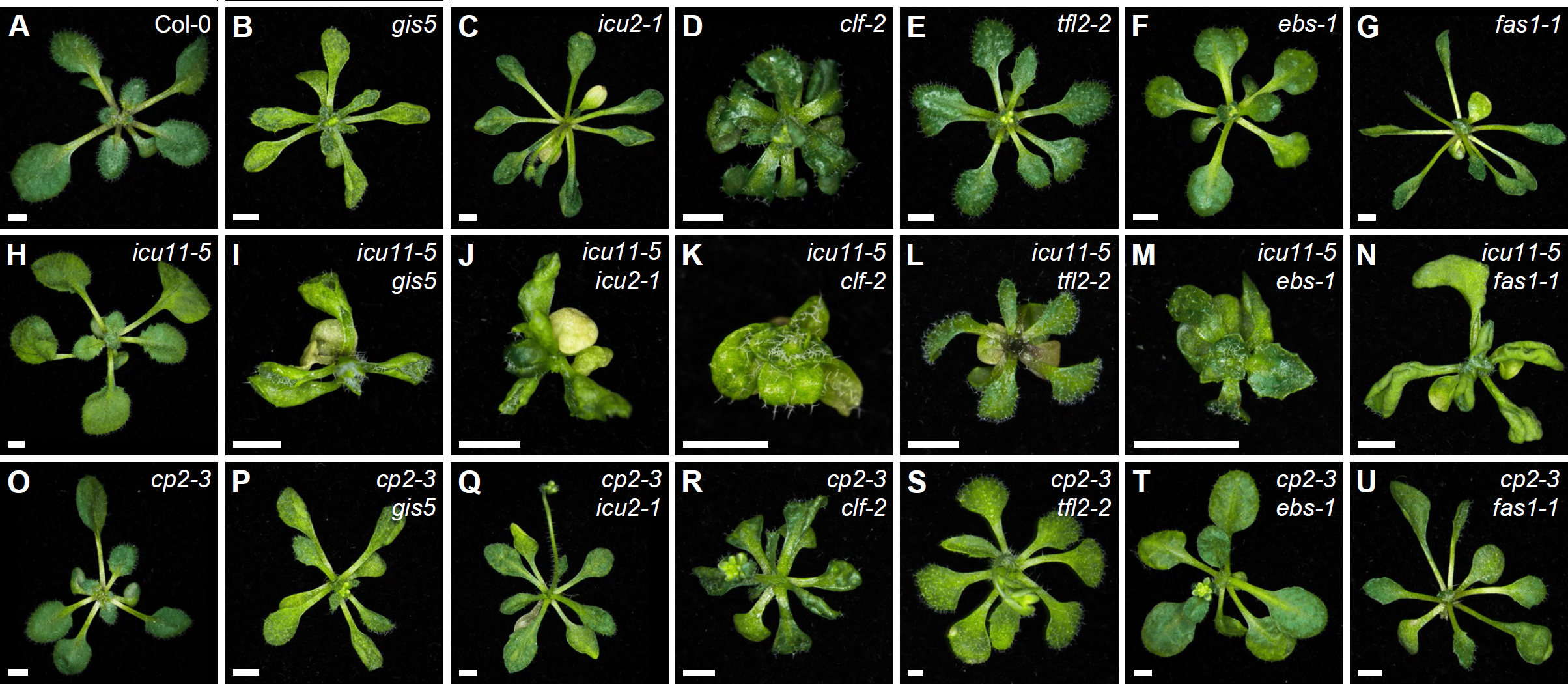
Phenotypes of double mutants involving *icu11-5* and *cp2-3* with *gis5*, *icu2-1*, *clf-2*, *tfl2-2*, *ebs-1*, and *fas1-1.* Rosettes of the wild-type Col-0 (A); the single mutants *gis5* (B), *icu2-1* (C), *clf-2* (D), *tfl2-2* (E), *ebs-1* (F), *fas1-1* (G), *icu11-5* (H), and *cp2-3* (O); and the double mutants *icu11-5 gis5* (I), *icu11-5 icu2-1* (J), *icu11-5 clf-2* (K), *icu11-5 tfl2-2* (L), *icu11-5 ebs-1* (M), *icu11-5 fas1-1* (N), *cp2-3 gis5* (P), *cp2-3 icu2-1* (Q), *cp2-3 clf-2* (R), *cp2-3 tfl2-2* (S), *cp2-3 ebs-1* (T), and *cp2-3 fas1-1* (U). Photographs were taken 15 das. Scale bars, 2 mm.

We also combined the mutations mentioned above with the *cp2-3* mutation, which lacks a distinctive phenotype; the resulting F_2_ progeny showed rosette phenotypes that were indistinguishable from those of their corresponding phenotypically mutant parent (Fig. 4P–U). We conclude that, unlike *ICU11*, the loss of *CP2* function is not sufficient to modify the phenotypes caused by mutations in other genes involved in epigenetic modifications, confirming the previously proposed unequal functional redundancy between *ICU11* and *CP2* (Mateo-Bonmatí et al., 2018).

### Sucrose partially rescues the lethality of the *icu11 cp2* double mutants

There is an expanding list of Arabidopsis mutants exhibiting morphological phenotypes that are partially or fully rescued by the exogenous supplementation of sucrose, such as *phosphatidylglycerolphosphate synthase 1* (*pgp1*) and *cyclophilin 38* (*cyp38*), which are defective in the assembly and proper function of photosystem II protein complexes, respectively (Fu et al., 2007; Kobayashi et al., 2016; Duan et al., 2021). This phenomenon is true for other Arabidopsis genes that are functionally unrelated, but whose mutations directly or indirectly impair or abolish photosynthesis. We observed the same effect in *icu11 cp2-1* and *icu11 cp2-2* double mutant seedlings, in which photosynthesis is diminished because they do not form true leaves, instead developing embryonic flowers immediately after germination. These mutants had a very slow growth rate, did not develop further, and died 20–40 days after stratification (das; Fig. 5A–C). When we raised the concentration of sucrose in the growth medium from 1% (w/v) to 3%, *icu11-5 cp2-1* double mutant plants developed main and axillary shoots with long internodes, small cauline leaves, disorganized flowers, and short siliques. Most of the disorganized flowers showed homeotic transformations of sepals and petals into carpels and had few stamens (Fig. 5D–H).

**Figure 5.**
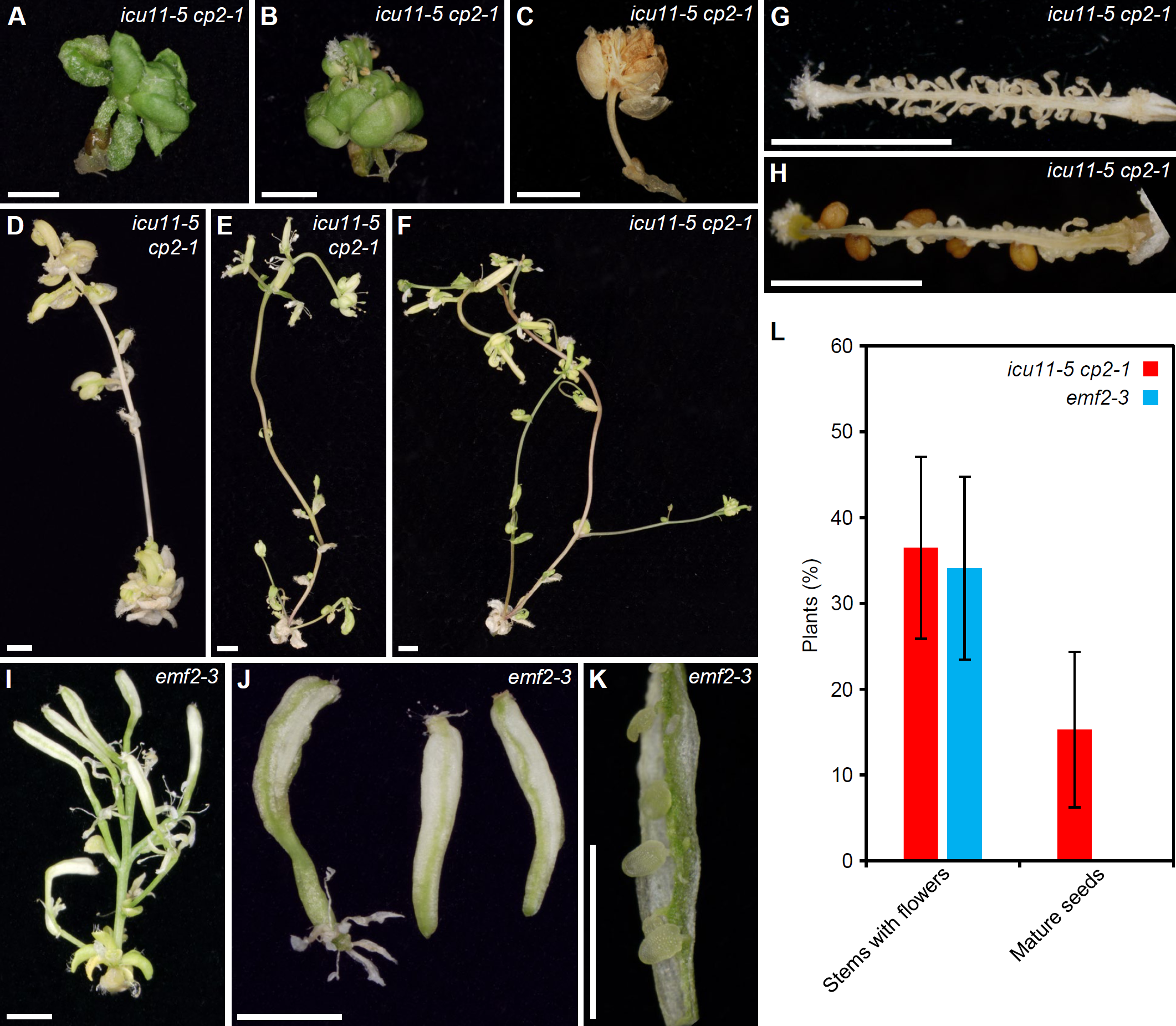
Partial rescue of the post-germinative lethality phenotype of *icu11-5 cp2-1* and *emf2-3* plants grown on culture medium supplemented with 3% sucrose. (A–K) Pictures of the *icu11-5 cp2-1* double mutant (A–H) or the *emf2-3* single mutant (I–K), showing embryonic flowers (A–C); seedlings developing stems (D–F), flowers (I), siliques (J), and dissected siliques (G, H, K). (L) Percentage of *icu11-5 cp2-1* and *emf2-3* plants with different phenotypes grown on culture medium supplemented with 3% sucrose. Error bars indicate standard deviation. A total of 136 *icu11-5 cp2-1* and 110 *emf2-3* plants were classified. Photographs were taken 26 (A, B), 36 (D–F, I–K), 60 (G, H), and 84 (C) das. Scale bars, 2 mm.

The *emf2-3* single mutant (Yoshida et al., 2001), whose morphological phenotype is similar to that of *icu11 cp2* double mutant plants, also exhibited a partial rescue under increased sucrose supplementation. These plants growing on 3% sucrose medium produced main and axillary shoots lacking apical dominance, as well as flowers with an altered structure that developed into short siliques with extended white sectors reminiscent of petal tissue (Fig. 5I–K). When grown on growth medium supplemented with 3% sucrose, 36.5% of *icu11-5 cp2-1* (*n* = 136) and 34.1% of *emf2-3* (n = 110) plants developed stems with flowers, although even with hand pollination only 15.3% of *icu11-5 cp2-1* plants and no *emf2-3* plants produced mature seeds (Fig. 5L). Only one out of the 23 seeds obtained in this manner from *icu11-5 cp2-1* plants germinated on medium supplemented with 3% sucrose, and still developed embryonic flowers.

An alteration in pollen viability may explain why only some *icu11-5 cp2-1* double mutant plants produced seeds. To assess pollen grain viability in the *icu11-5*, *cp2-1*, and *icu11-5 cp2-1* mutants, we stained their anthers with Alexander solution. We detected no aberrations in anther shape or size or in pollen grain viability for the *cp2-1* and *icu11-5* anthers when compared to Col-0 (Fig. 6A–F); however, 66% of the observed *icu11-5 cp2-1* anthers were smaller and carried fewer but viable pollen grains (Fig. 6G, H, K). The remaining 34% of anthers were also small and contained only non-viable pollen that turned blue/gray upon staining (Fig. 6G–K).

**Figure 6.**
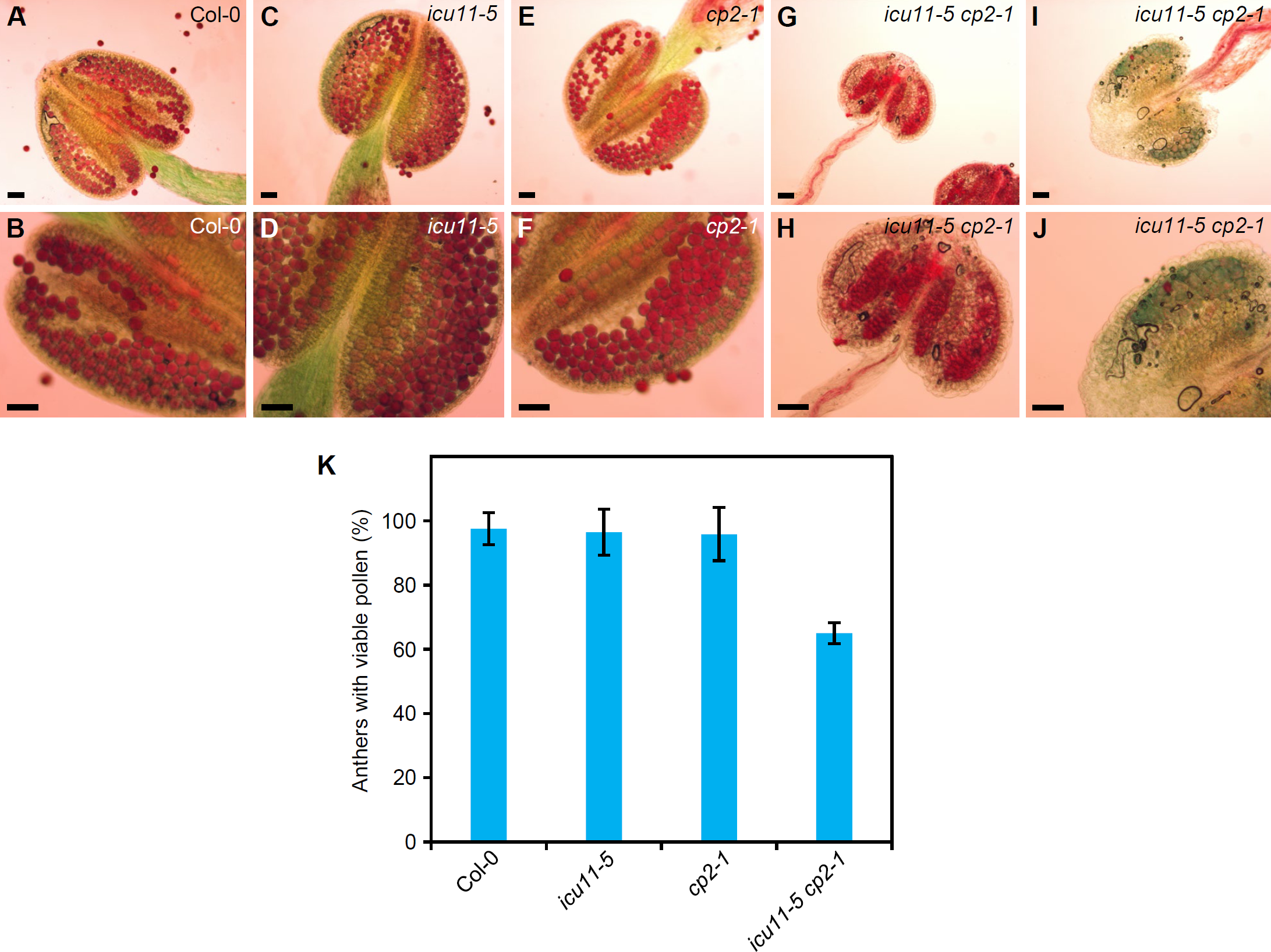
Pollen grain viability in *icu11-5*, *cp2-1*, and *icu11-5 cp2-1*. (A–J) Anthers of Col-0 (A–B), *icu11-5* (C–D), *cp2-1* (E–F), and *icu11-5 cp2-1* plants (G–J). Col-0, *icu11-5*, and *cp2-1* plants were grown on standard culture medium, while *icu11-5 cp2-1* plants were grown on medium supplemented with 3% sucrose. Purple/red and blue/gray grains are viable and non-viable pollen, respectively. All samples were treated for 5 min with Alexander solution. Scale bars, 50 μm. (K) Percentage of anthers containing viable pollen after collecting and studying 28 anthers (three or four per plant). Error bars indicate standard deviation.

## DISCUSSION

### The severity of leaf aberrations in the *icu11* single mutants is dependent on genetic background

An analysis of phenotypic differences both between and within different single mutant alleles, as well as a comparison to their corresponding double mutants, is a common task in developmental genetics. Many studies have shown that morphological phenotypes can vary depending on the genetic background; for example, the *gibberellin biosynthesis* (*ga5*) mutant in L*er* displays a much more pronounced drop in shoot fresh weight than the *ga20ox1-3* (another *ga5* loss-of-function allele) mutant, in the Col-0 background (Barboza-Barquero et al., 2015). Another example is the introduction of either the wild-type L*er* allele or the loss-of-function allele from the Japanese accession Fuk for the MADS box transcription factor gene *SHORT VEGETATIVE PHASE* (*SVP*), which regulates flowering time, into five Arabidopsis accessions. In some of these accessions, the presence of *SVP*-Fuk accelerated flowering, whereas *SVP*-L*er* delayed it; however, in other accessions, no noticeable differences were noted (Méndez-Vigo et al., 2013).

The *icu11-1* mutant exhibits a developmental phenotype previously observed in other mutants that carry alleles of genes encoding components of the epigenetic machinery (Mateo-Bonmatí et al., 2018). These characteristic aberrations, such as leaf hyponasty and early flowering, are in some cases associated with a reduced deposition of the epigenetic mark H3K27me3 across a large number of genes (Förderer et al., 2016). The gene-edited *icu11-5* and *icu11-6* null alleles of *ICU11* that we obtained in the Col-0 genetic background in this study exhibited a leaf incurvature milder than that of *icu11-1* in the S96 background. These differences are most likely due to differences in the genetic backgrounds, as we also obtained two CRISPR/Cas9 mutants in the S96 background, *icu11-4* and *icu11-7*, that displayed identical leaf curvature as *icu11-1.* Furthermore, it is worth mentioning here that *icu11-6* and *icu11-7* carry independently obtained but identical mutations: a deletion of one nucleotide immediately downstream of the PAM; therefore, their differences in leaf phenotype can only be due to their different genetic backgrounds.

### The unequal functional redundancy between the *ICU11* and *CP2* paralogs is not dependent on genetic background

*CP2* and *ICU11* are unequally redundant paralogs, as we inferred from the phenotype of the *icu11-1/icu11-1;CP2/cp2-3* sesquimutant, which develops lethal embryonic flowers, while the reciprocal *ICU11/icu11-1;cp2-3/cp2-3* sesquimutant is phenotypically wild type (Mateo-Bonmatí et al., 2018). The *icu11-1 cp2-1* and *icu11-1 cp2-2* double mutants also developed embryonic flowers, whereas the *ICU11/icu11-1;cp2-1/cp2-1* and *ICU11/icu11-1;cp2-2/cp2-2* sesquimutants were indistinguishable from wild type plants, and the *icu11-1/icu11-1;CP2/cp2-1* and *icu11-1/icu11-1;CP2/cp2-2* sesquimutants were indistinguishable from *icu11-1* single mutant plants. While the genetic background of these double mutants and sesquimutants was hybrid (S96/Col-0), all combinations between the *icu11-5*, *icu11-6*, *cp2-1*, *cp2-2*, and *cp2-3* mutations that we obtained here were in a single genetic background (Col-0). Furthermore, as previously shown for *icu11-1/icu11-1;cp2-3/cp2-3*, no *icu11-5/icu11-5;cp2-3/cp2-3* or *icu11-6/icu11-6;cp2-3/cp2-3* double mutants were obtained likely because of their gametic or early embryonic mortality. The genetic combination of our new *icu11* alleles with *cp2* alleles, all in the Col-0 background, confirmed that *CP2* behaves as a haploinsufficient locus in a homozygous *icu11* background (Pérez-Pérez et al., 2009; Meinke, 2013), and in particular, that the embryonic flower phenotype of the double mutant and some sesquimutant combinations of their alleles are not influenced by the S96, Ws-2, or Col-0 genetic backgrounds.

We also provided further evidence for the unequal functional redundancy between *ICU11* and *CP2* through the analysis of their genetic interactions with loss-of-function alleles of other epigenetic machinery components. We established that *icu11-5* synergistically interacts with *clf-2*, *ebs-1*, *gis5*, *icu2-1*, *tfl2-2*, and *fas1-1* in the presence of two wild-type *CP2* copies, as previously reported for *icu11-1*. There were some minor differences between the synergistic phenotypes of the double mutant combinations of *ebs-1* (in the L*er* background), *tfl2-2* (Col-0), and *fas1-1* (En-2) with *icu11-1* (S96) or *icu11-5* (Col-0), which can be attributed to the genetic background. By contrast, the double mutant combinations of *ebs-1*, *tfl2-2*, or *fas1-1* with the *cp2-3* null allele, in the presence of two *ICU11* wild-type copies, resulted in phenotypes indistinguishable from those of the *ebs-1*, *tfl2-2*, or *fas1-1* single mutants. Apparently, the presence of a wild-type allele of *ICU11* impedes the identification of *CP2* genetic interactors, as they are redundant. Thus, the lethality of the *icu11 cp2* double mutant is an obstacle for an independent characterization of *CP2*.

### The viability of the *icu11 cp2* double mutants and sesquimutants is dependent on exogenous carbon

The phenotype of a large number of mutants in genes that are primarily associated with photosynthesis or processes closely linked to it can be partially or completely alleviated by supplementation with an exogenous carbon source. Indeed, photosynthesis, primarily occurring in plant leaves, serves as the ultimate source of sugars, which are the primary carriers of both sunlight energy and carbon required for metabolism. Sugar availability is crucial for Arabidopsis development, leading to early developmental arrest in mutants with impaired photosynthesis, such as *pgp1* (Kobayashi et al., 2016), and *fructokinase-like 2-4* (*fln2-4*; Huang et al., 2013) or *white cotyledons* (*wco*; Yamamoto et al., 2000). Since *WCO* is only required for chloroplast biogenesis in cotyledons but not in true leaves, *wco* mutants can be supplied with 3% sucrose for a few days to enable them to survive and produce true leaves, after which the plants develop normally. Other mutations of genes involved in photosynthesis are not seedling or plant lethal but delay growth; for example, CYP38 participates in the assembly and maintenance of photosystem II, and loss-of-function *cyp38* alleles show retarded growth and pale green leaves. A 1% sucrose supplementation was sufficient to rescue rosette growth, although *cyp38* still displayed hypersensitivity to light, while higher concentrations of sucrose increased the length of the primary root (Fu et al., 2007; Duan et al., 2021).

Although several genes repress flowering in Arabidopsis, the *emf* single mutants are unique because they completely skip the vegetative phase after seed germination without generating true leaves (Sung et al., 1992; Yang et al., 1995). The *icu11 cp2* double mutants resembled the *emf* single mutants, as they also lacked true leaves. Impaired carbon fixation results in the early-development arrest of these mutants, so they cannot produce reproductive structures beyond a disorganized embryonic flower. These mutants represent a class completely different from those mentioned above, as they are not primarily affected in genes related to photosynthesis. Instead, they skip vegetative development and die because they cannot develop leaves, which does not allow them to photosynthesize properly.

### ICU11 and CP2 appear to regulate flower organ identity genes

The increase from 1% to 3% sucrose in the growth medium was sufficient to allow the formation of reproductive-phase structures in our *icu11 cp2* double mutants such as axillary shoots, cauline leaves, and disorganized inflorescences exhibiting homeotic transformations. Analysis of these structures revealed that *ICU11* and *CP2* are not only required for vegetative development, as previously described (Mateo-Bonmatí et al., 2018), but also ensure proper reproductive development. *ICU11* and *CP2* are expressed in both vegetative and reproductive organs, with *CP2* showing higher expression levels than *ICU11* in the flowers and siliques of wild-type plants (Mateo-Bonmatí et al., 2018). Homeotic transformations have been described as a consequence of the overexpression or loss-of-function of the floral organ identity genes *APETALA1* (*AP1*), *AP2*, *AP3*, *PISTILLATA* (*PI*), *AGAMOUS* (*AG*), *SEPALLATA1* (*SEP1*), *SEP2*, and *SEP3* (Wellmer et al., 2014; Goslin et al., 2023).

Similar to homeotic transformations in *icu11-5 cp2-1*, the loss-of-function *ap2* mutations and the overexpression of *AG* or *SEP3* (classes C and E of the ABC model of floral development, respectively) result in the transformation of sepals and petals into carpelloid structures (Mizukami and Ma, 1992; Riechmann and Meyerowitz, 1997; Chen, 2004; Castillejo et al., 2005). Moreover, the weak *emf2-10* allele causes the appearance of carpelloid sepals (Chanvivattana et al., 2004) reminiscent of those of the *icu11-5 cp2-1* double mutant. Indeed, previous studies have described that ICU11 plays a role in repressing *AG*, *SEP1*, *SEP2*, and *SEP3* expression during the vegetative phase (Mateo-Bonmatí et al., 2018; Bloomer et al., 2020). Therefore, based on our observations of *icu11-5 cp2-1*, we propose that ICU11 and CP2 regulate the expression of floral identity genes during reproductive development.

The homeotic transformations of the *icu11-5 cp2-1* and *emf2-3* flowers impeded self-pollination. Hand pollination of the *icu11-5 cp2-1* double mutant, but not the *emf2-3* single mutant, resulted in a few seeds, only one of which germinated and developed an embryonic flower when grown on growth medium containing 3% sucrose. In this growth condition, *icu11-5 cp2-1* anthers were smaller than those of Col-0, and only 64% contained small but viable pollen grains. When grown under short-day conditions, the mutant *emf2-3* also develops a small shoot with two or three flowers and short siliques that did not produce mature seeds (Chen et al., 1997).

## ACKNOWLEDGMENTS

The authors wish to thank J.M. Serrano and J. Castelló for their excellent technical assistance. Research in the laboratory of J.L.M. was supported by grants from the Ministerio de Ciencia e Innovación of Spain (PGC2018-093445-B-I00, EQC2018-005181-P and EQC2019-006592-P [MCI/AEI/FEDER, UE]) and the Generalitat Valenciana (PROMETEO/2019/117 and IDIFEDER/2020/019). R.N., E.M.-B., and L.J.-V. held predoctoral fellowships from the Generalitat Valenciana (GRISOLIAP/2016/131) and the Ministerio de Universidades of Spain (FPU13/00371 and FPU16/03772), respectively.

## AUTHOR CONTRIBUTIONS

J.L.M. conceived and supervised the study, provided resources, and obtained funding. R.N., L.J.-V., and J.L.M. designed the methodology. R.N., L.J.-V., and E.M.-B. performed the experiments. R.N., L.J.-V., and J.L.M. wrote the original draft. All authors reviewed and edited the manuscript.

## SUPPLEMENTARY MATERIAL

**Supplementary Figure S1**. Design and effects of the CRISPR/Cas9 mutagenesis of *ICU11*.

**Supplementary Figure S2**. Testing of two putative CRISPR/Cas9 off-targets in new *icu11* mutant lines.

**Supplementary Table S1**. Primer sets used in this work.

**Supplementary Table S2**. Double mutant combinations that rendered additive phenotypes, obtained by crossing *icu11-1* to mutants carrying alleles of genes encoding known plant epigenetic machinery components.

