## Supplementary Figures and Table S1 for "The unequal functional redundancy of Arabidopsis *INCURVATA11* and *CUPULIFORMIS2* is not dependent on genetic background"

### Supplementary Figures and Tables

Supplementary Material included in this file:

- Supplementary Figures S1 and S2
- Supplementary Table S1
- Supplementary References

Supplementary Material not included in this file:

- Supplementary Table S2

**A** Features of sgRNA and its target

sgRNA name: ICU11_sgRNA1

sgRNA sequence: GCGAGGCAAGATTGAAGCTTCGG

Target location: Chr1:8127046. AT1G22950, first exon

Target strand: complementary

sgRNA efficiency: 49.72%

sgRNA predicted off-targets: MM(0):0, MM(1):0, MM(2):0, MM(3):0, MM(4):2

**B**

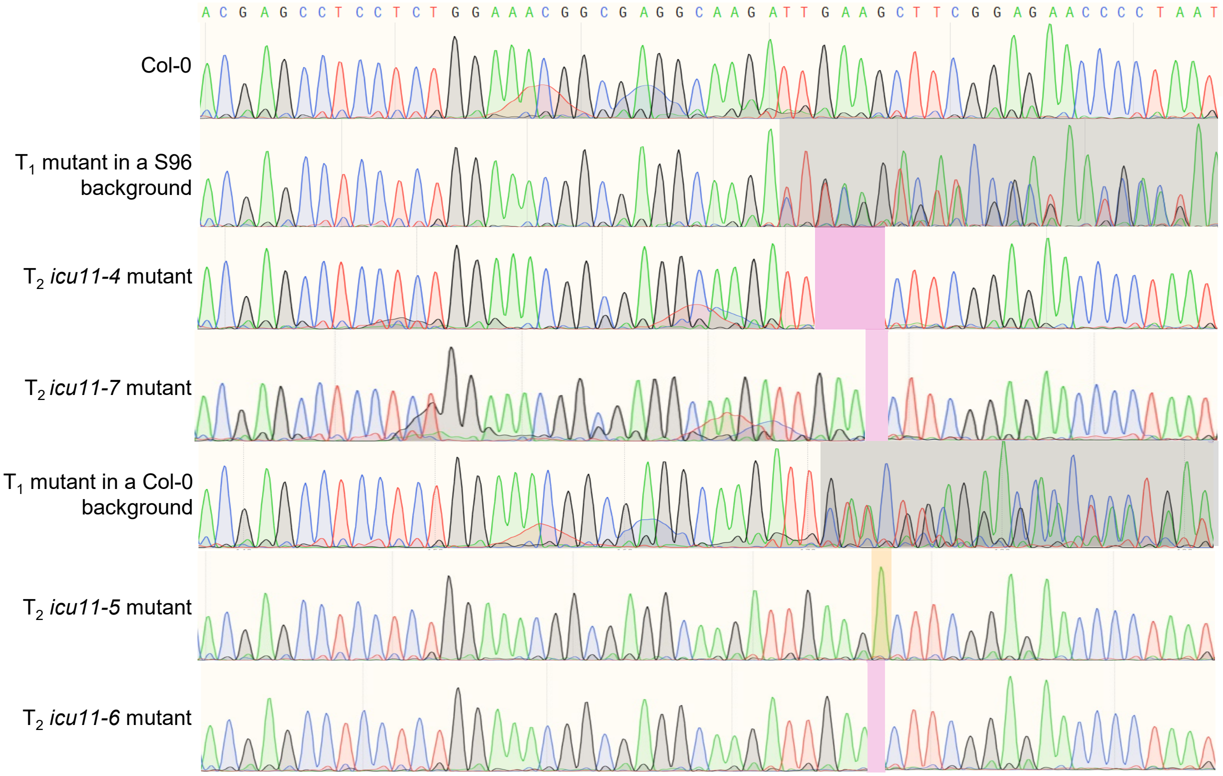


**C**

```
ICU11-4      1 MCNQTPLRSMALDSSGKQPEQQQQQQPRASSGNGEARFFGEPLMKNMSLRTMKIYLWTIV
ICU11-6      1 MCNQTPLRSMALDSSGKQPEQQQQQQPRASSGNGEARLNFGEPLMKNMSLRTMKIYLWTI
ICU11-7      1 MCNQTPLRSMALDSSGKQPEQQQQQQPRASSGNGEARLNFGEPLMKNMSLRTMKIYLWTI
ICU11-5      1 MCNQTPLRSMALDSSGKQPEQQQQQQPRASSGNGEARLKTSENP----------------
ICU11        1 MCNQTPLRSMALDSSGKQPEQQQQQQPRASSGNGEARLKLRRTPNEEHEPENYEDLPLDY

ICU11-4     61 LLCSPLLSVTYLSSFSIPHESIKLVS----------------------------------
ICU11-6     61 VLLCSPLLSVTYLSSFSIPHESIKLVS---------------------------------
ICU11-7     61 VLLCSPLLSVTYLSSFSIPHESIKLVS---------------------------------
ICU11-5        ------------------------------------------------------------
ICU11       61 SPSLFTSLERYLPEQLLNSTRIDKASFMRDLLLRYSPDTERVRVLRHKEYRDKIMSSYQR
```

**Supplementary Figure S1.** Design and effects of the CRISPR/Cas9 mutagenesis of *ICU11*. (A) Details of the *ICU11* sgRNA1 target. The PAM sequence is shown in red. On-target mutation efficiency was calculated using the "Rule Set 2" scoring model, which provides values ranging from 0 to 100 (Doench et al., 2016). Possible off-target events are represented according to the number of mismatches [MM (number)]. (B) Electropherograms of the *ICU11* sgRNA1 target site in wild-type plants, and $T_1$ and $T_2$ transgenic plants. The gray, magenta, and orange shaded areas indicate chimeric, deletion, and insertion mutations, respectively. (C) Multiple amino acid sequence alignment of the predicted proteins translated from wild-type and CRISPR/Cas9 alleles showing that all the latter produce truncated proteins. Identical and similar residues are shaded in black and gray, respectively. Numbers indicate residue positions.

**A** Details of two putative off-targets of ICU11_sgRNA1

| | ICU11_sgRNA1 off-target 1 | ICU11_sgRNA1 off-target 2 |
|---|---|---|
| Chromosome | 1 | 3 |
| Position | 19132845 | 2280675 |
| Strand | Forward | Complementary |
| Gene | Intergenic, between AT1G51590 (*MNS1*) and AT1G51600 (*GATA28*) | First exon of AT3G07170 (*IRP1*) |
| Mismatches | ICU11-T1: **GCGAGGCAAGATTGAAGCTTCGG**<br>Col-0: **GCGTGGAAATATTGAAACTTTGG** | ICU11-T1: **GCGAGGCAAGATTGAAGCTTCGG**<br>Col-0: **GCGAAGCAAGAATCAAACTTGGG** |

**B**

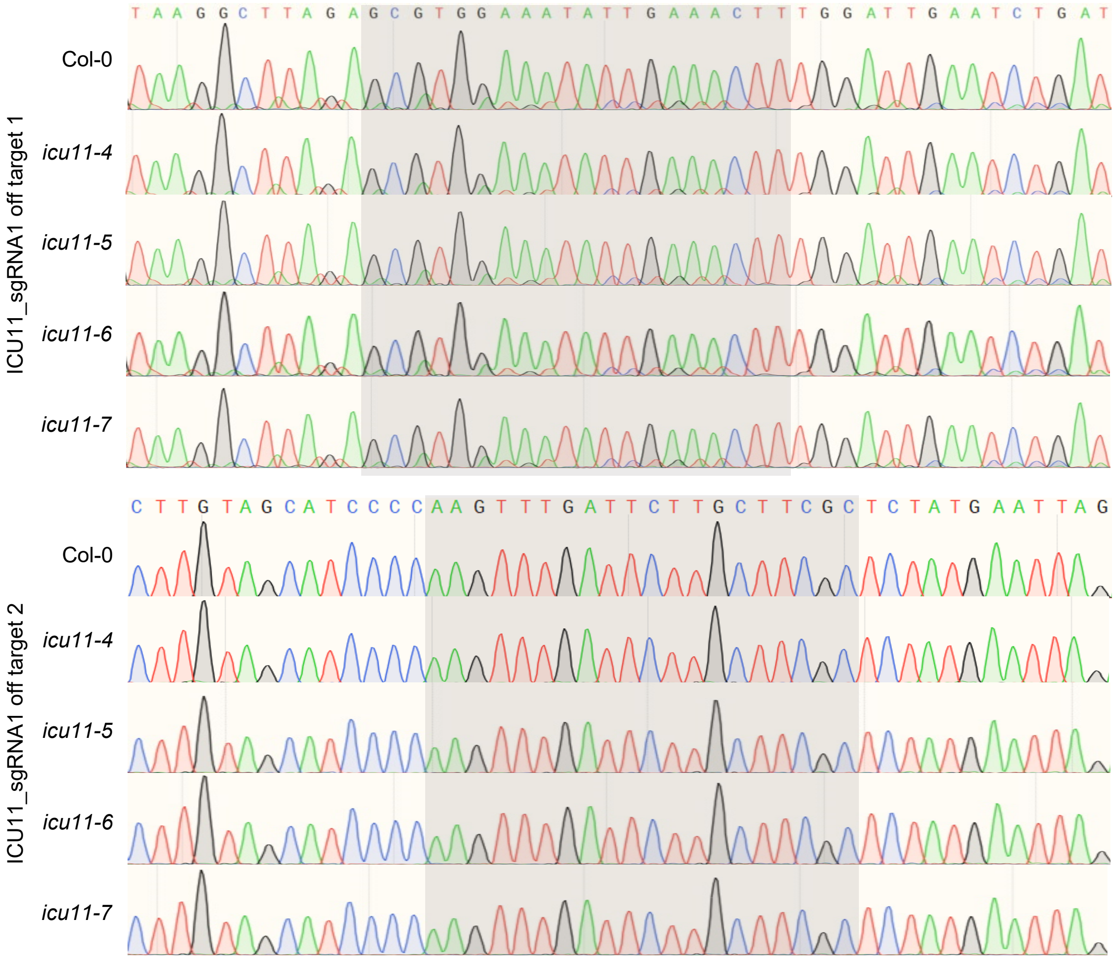


**Supplementary Figure S2.** Testing of two putative CRISPR/Cas9 off-targets in the new *icu11* mutant lines. (A) Off-targets were found using Cas-OFFinder, a bioinformatic tool developed by Bae et al. (2014). Mismatches are shown in red, with the first nucleotide of the PAM sequence (NGG) is in blue. (B) Sanger sequencing electropherograms of putative off-targets in $T_3$ mutant and wild-type plants. The gray shaded area corresponds to the putative off-target sequence.

**Supplementary Table S1.** Primer sets used in this work

| Purpose | Oligonucleotide name(s) | Oligonucleotide sequences (5′ → 3′) | |
|---|---|---|---|
| | | Forward primer (F) | Reverse primer (R) |
| Genotyping | ICU11-Off-target1_F/R | TGGTGGGTTTGGTTTGTCTC | CTCGGTCATTGGAGCAACTT |
| | ICU11-Off-target2_F/R | TGAGTCTGGAAGCAGGAAGG | AATGGGCAAATCAGAGAGTCC |
| | At1g22950_1F/R | ACCCTAACCTCTCAAACAAACCA | AGACTTTGTTAACCCAATCCGAC |
| | At1g22950_4F/R | CCTCTCAAACAAACCATCATCA | CGCTCAGTATCAGGGGAATATC |
| | SAIL_1215_B02_L/R | GAGCGATAACAGTGAGCTTGG | GACATTTTCAAACCATTCATGC |
| | SAIL_658_E12_L/R | AGAGGCAAGAGACGAAAAAGC | CCTTTGAGCCTGTAGCATCAG |
| | SAIL_621_G08_L/R | TGAGAGCGAAAGCTTTCATTC | AACAAATGACTGGAGCAGAGC |
| | gis-5_F/R | GAAGCAAGAACAGGTTTCTATG | AGCTAGTTACACTCGAGGATA |
| | icu2-1_F/R | TGTTGAAGGAGGTCAGTTATTCT | CACAAGTGTTTTGGATGACTGAA |
| | clf-2_F/R | ATGGCGTCAGAAGCTTCGCC | CTGGACCTCTCTCCTCCGC |
| | tfl2-2_F/R | TATCAGCGGTGATCGGTGTG | CGCCGTAATTCTCCCGGTAA |
| | ebs-1_F/R | TGAAGGTGTGAACAATGCAT | GAAAACTCGACCTGGTGTCG |
| | fas1-1_F/R | TGAGCTGTTCTTCTGCATCATG | ACTATGGTAGCTGTGAAGAGTG |
| | AT5G51230_1F/R | TGTAATGGTTCAGAGATCAATAGAA | GTCCGTGCAATCTTGAGAATG |
| | SALK_131712_L/R | CAGAAGAAGATCGTCCGAGTG | TGAACTTCCCCACTCTTCATG |
| | SALK_056440_L/R | TGGTCAGATGGGCTAGAATTG | AACGCGTTGCTGTAGAAACTC |
| | SAIL_826_A06_L/R | AGCAGCAGAAGAAGAAGCATG | TTTGGCCTACAAAGACACCAG |
| | SALK_021316_L/R | GAGCCGTCTCATCAAACTGAC | TTGCAGGAGCAAATATGGAAC |
| | SALK_150863_L/R | AGATCGCTTCCAGAGTTAGCC | TTGTCGCAAAAAGCAAAAGAG |
| | SAIL_223_F05_L/R | GGATCAGCCAAAAGGTTAAGG | TCATTCACTTTGCATCACTCG |
| | SAIL_809_E03_L/R | GCGTGTACCAGTTTCAAGGAG | TAAAGAGCCCAGTTGTGAAGC |
| | SALK_045303_L/R | CCAGTTAAGGACAGAACACCG | TCGTCTTTCGATCAAATCCAC |
| | SALK_022363_L/R | ATCAATGTGGCATCTAGTGGC | ACCCGCCTCTTCTTCATCTAC |

**Supplementary Table S1 (continued).** Primer sets used in this work

| Purpose | Oligonucleotide name(s) | Oligonucleotide sequences (5′ → 3′) | |
|---|---|---|---|
| | | Forward primer (L or F) | Reverse primer (R) |
| Genotyping | SAIL_97_E06_L/R | CTTTCCCAGTTTTTACTGCCC | AATCACTCGCTTCTTCCACTG |
| | SALK_149002_L/R | AATGAAAGCATGCGGATACAC | TCCGTGTTGACTGGAAAGATC |
| | SALK_130607_L/R | TTTCTCTTGTCCGGTGAAATG | CCTGCAACAATCAGTGTGATG |
| | SAIL_240_H01_L/R | TTGAGATGAATCTGGAGACCG | AAACGACGACGTATTGGAGTG |
| | SALK_149692_L/R | TCTTGTGACAGGTGCAACTTG | AAACAAAGCTAGGCACAAGGC |
| | SALK_080380_L/R | AGGGAACATGTCATCCATGAG | AGGGAGAATCTGAGAACCTGC |
| | SALK_027726_L/R | ATGGTGTGCGAATCTATGACC | ACGGAGAGGAAAGCTCAAGAC |
| | LB1[1] | GCCTTTTCAGAAATGGATAAATAGCCTTGCTTCC | |
| | LbB1.3[2] | ATTTTGCCGATTTCGGAAC | |
| Cloning | ICU11_sgRNA1_F/R | ATTGCGAGGCAAGATTGAAGCTT | AAACAAGCTTCAATCTTGCCTCGC |

[1,2]These primers were used for genotyping [1]SAIL and [2]SALK lines, and their sequences were taken from [1]Sessions et al. (2002) and [2]T-DNA Primer Design (http://signal.salk.edu/tdnaprimers.2.html).
